# The dorsal striatum sets the sensitivity to effort

**DOI:** 10.1101/2020.03.13.991000

**Authors:** Maria-Teresa Jurado-Parras, Mostafa Safaie, Stefania Sarno, Jordane Louis, Corane Karoutchi, Bastien Berret, David Robbe

## Abstract

The dorsal striatum (dS) has been implicated in storing and retrieving procedural memories and controlling movement kinematics (e.g., speed). Since procedural memories are expressed through movements, the exact nature of the dS function has proven difficult to delineate. Here we challenged rats in complementary tasks designed to alleviate this performance confound. Surprisingly, dS lesions spared task-specific procedural memories but altered the kinematics of their expression in motor routines. Further behavioral analyses combined with simulations in the optimal control framework indicated that these alterations reflected an increased sensitivity to effort with preserved reward-seeking and ability to modulate movement speed. By setting the sensitivity to effort, the dS contributes to the optimization of the energy invested into voluntary movements. Such an elementary function of the dS might explain its implication in both procedural decisions and the control of movement speed.

## Main Text

The dorsal striatum (dS) plays a critical role in the control of voluntary actions. Activation of dS projection neurons forming the direct (indirect) basal ganglia pathway facilitates (prevents) movement production through disinhibition (inhibition) of brainstem and forebrain motor regions [1, 2]. This fundamental feature of the basal ganglia’s functional anatomy, combined with recordings and perturbations of neuronal activity in various behavioral tasks, has led to two main hypotheses regarding the nature of the dS motor function: 1) a role in learning and performing action sequences, which are also sometimes referred to as procedural skills or memories [3, 4, 5, 6, 7, 8, 9, 10, 11]; 2) a role in controlling movement speed [2, 12, 13, 14, 15, 16]. Testing the relative validity of these hypotheses is challenging because movements are typically used as a readout of procedural memories. Consequently, in procedural learning tasks used to probe the dS function through perturbation of neuronal activity, it is nearly impossible to disentangle whether impaired performance is due to inability to implement a correct procedural decision into movements, or an impaired retrieval of a procedural memory with preserved motor control ability.

To understand how the dS contributes to the control of voluntary actions while limiting as much as possible the above performance confound, we challenged rats in a set of motor tasks taking place on a motorized treadmill. In the first task [17], to obtain a drop of sucrose solution, rats had to wait for a fixed goal time (*GT* = 7 s relative to treadmill onset) before entering a reward area located at the front of the treadmill, while its belt was slowly moving backward (Fig. 1A). Across practice sessions composed of ∼130 trials (treadmill on) interleaved with intertrials (treadmill off), animals learned to wait longer and longer before entering the reward area, until reliably doing it very close to the *GT* (Fig. 1B, Fig. S1). Task proficiency was clearly associated with the acquisition and reliable performance of the following routine (Fig. 1C, compare left and middle): 1) during the intertrial, following the consumption of sweetened water, rats remained in the reward area; 2) when the treadmill was turned on (trial onset), they did nothing and let the moving belt carry them away from the reward area; 3) when they reached the rear wall of the treadmill, they outran the opposing treadmill to re-enter the reward area just after 7 s (Entrance Time, *ET* > *GT*). After 2-3 weeks of daily practice, rats used this wait-and-run routine in about 75% of the trials (Fig. 1C, right). Finally, learning this routine was paralleled by a robust invigoration of the running phase toward the reward area (i.e., during the first 10 sessions, rats crossed the treadmill toward the reward area with progressively faster speeds, Fig. 1D). Rats could have used several other strategies to perform proficiently in this task. For instance, they could have remained close to the reward area by running at the same speed as the treadmill for several seconds, and then perform a short acceleration to enter the reward area just after *GT*. Still, we recently reported that the usage of the wait-and-run routine facilitated timing accuracy and that a majority of animals relied on this strategy [17], despite the fact it might not be the optimal solution in terms of effort.

**Figure 1:**
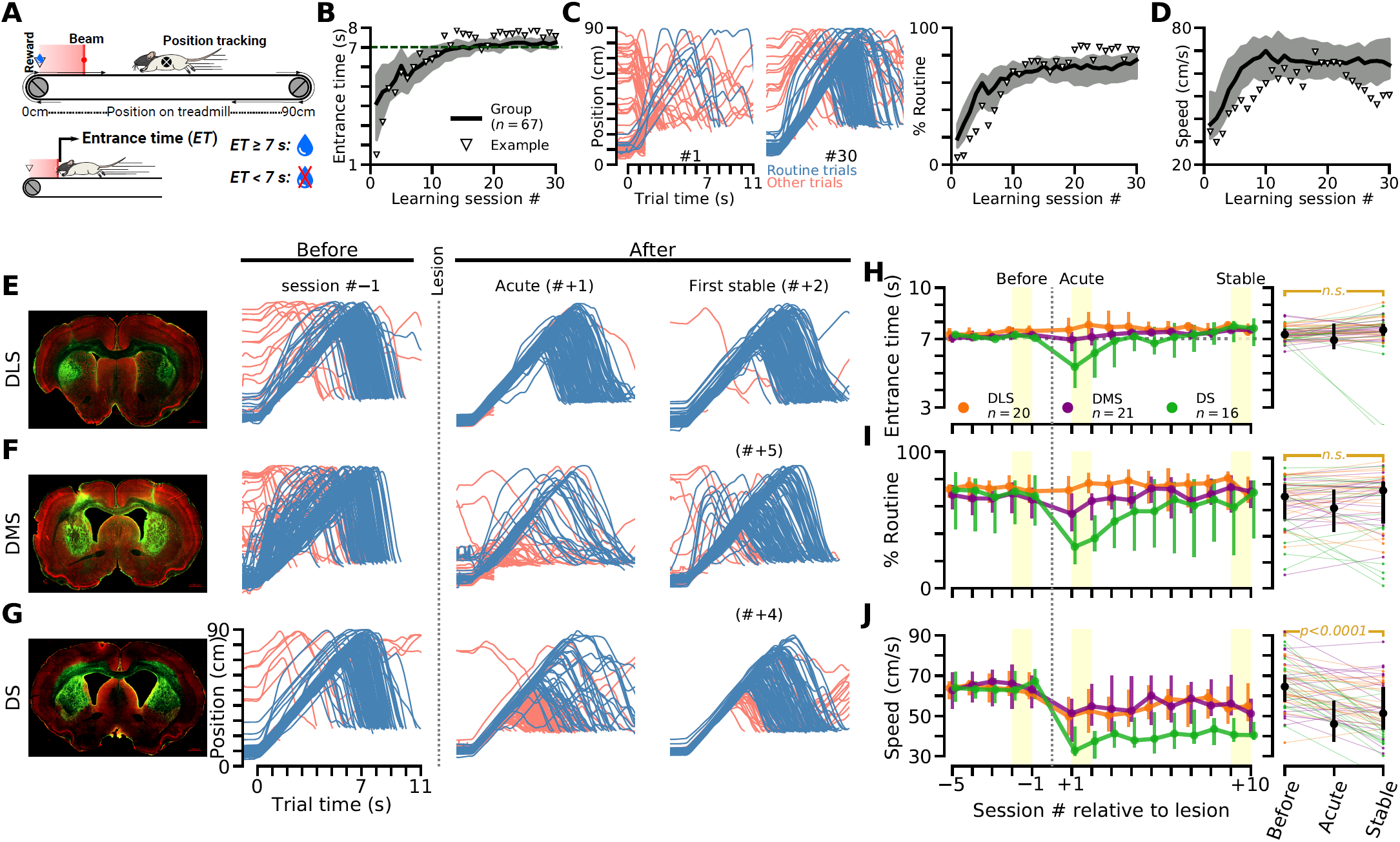
The dorsal striatum is necessary to invigorate the running component of a motor routine. **(A)** Experimental apparatus and task rules. **(B)** Entrance time across training sessions for all the rats trained in this task. Shaded area represents the interquartile range. **(C)** Trajectories of an example animal on the treadmill, for all the trials performed during sessions #1 and #30 (left). Percentage of trials during which animals performed the wait-and-run routine, across sessions (right). **(D)** Running speed when animals ran toward the reward area, across sessions. Triangles in B to D indicate the changes in performance for the example animal whose trajectories are shown in C (left). **(E-G)** Histology (1^*st*^ column, GFAP in green shows gliosis, red is NeuN) and trajectories of single animals with bilateral lesions of the dorsolateral, dorsomedial and dorsal striatum (E: DLS, F: DMS: G: DS). # indicates session number relative to lesion break. **(H-J)** Left, Time course of the lesion effect on ET (H), percentage of routine usage (I) and running speed (I). Right, group data statistical comparison before vs after lesion (10,000 resamples). Trajectories in C, E, F and G are cut after ET.

We then performed fiber-sparing lesions of the dS in 57 animals well-trained in this task. The lesions targeted either the dorsolateral or the dorsomedial region of the striatum or both territories (DLS, DMS, DS lesions, Fig. 1, E to G, Fig. S2). Behavioral testing resumed two weeks after the lesion surgery (Fig. S3). The average *ET* of animals with the largest lesions (mostly DS lesions) dropped during the first post-lesion session (Fig. 1, F to H). These animals ran toward the reward area prematurely after trial onset (Fig. 1, F to H) and, consequently, a reduction in the usage of the wait-and-run routine was observed during this first post-lesion session (Fig. 1, F, G and I). But surprisingly, most of these animals recovered from this initial impairment and after a few additional sessions, task proficiency was similar to the pre-lesion level (Fig. 1, H and I). Moreover, for most of the animals with a lesion restricted to the DLS or DMS, task proficiency was virtually unaltered (Fig. 1, E, H and I). In contrast, the animals’ speed when they ran toward the reward area was irreversibly decreased following lesion (Fig. 1, E to G and J), an effect that robustly correlated with lesion size (Fig. S4). Noticeably, the maintained task proficiency following dS lesion suggested that the motivation of the animals to obtain rewards (reward-seeking) was preserved. Accordingly, animals with a dS lesion kept licking the sweetened water upon correct *ET*, although licking initiation was delayed (Fig. S5).

These results suggest that the dS is selectively critical for the invigoration of the reward-oriented active component of the wait-and-run routine. But at this stage, we cannot rule out that the transient impairment in performance induced by large dS lesions is caused by a deterioration of procedural memory and reflects a reversal to pre-learning behavior (Fig. 1, B and C left, [11]), compensated in subsequent post-lesion sessions through a dS-independent learning process. We thus trained a new group of animals in a modified version of the task in which the treadmill belt moved slowly toward the reward area (instead of away from it). Several animals learned to perform proficiently this version of the task by adopting a *run-and-wait* routine: they moved to the back of the treadmill during intertrial (after licking the sweetened water from the previous trial and while the belt was immobile) and, following trial onset, they remained immobile while being passively transported toward the reward area (Fig. 2, A to C). Regarding the aforementioned issue of performance confound, a key feature of this routine is that it can be performed even with a compromised motor system. Indeed, to be proficient in this task, animals just have to reach the back of the treadmill during the relatively long (20 s) intertrial. Thus, a deficit in performing this routine would most likely reflect an impaired procedural memory. Concretely, if the dS lesion abolished the procedural memory underlying the run-and-wait routine, we expected that, in the first post-lesion session, animals started the trials close to the reward area, as they spontaneously did before learning. This is not what was observed and, generally, task proficiency was preserved following dS lesion (Fig. 2, B to D). A lack of effect of the dS lesion was not due to the fact that learning the run-and-wait routine was easier than learning the wait-and-run routine (Fig. S6). Thus, the transiently impaired performance of the wait-and-run routine, seen in animals with a large dS lesion (Fig. 1, F to I), is unlikely to be caused by a deterioration of procedural memory.

**Figure 2:**
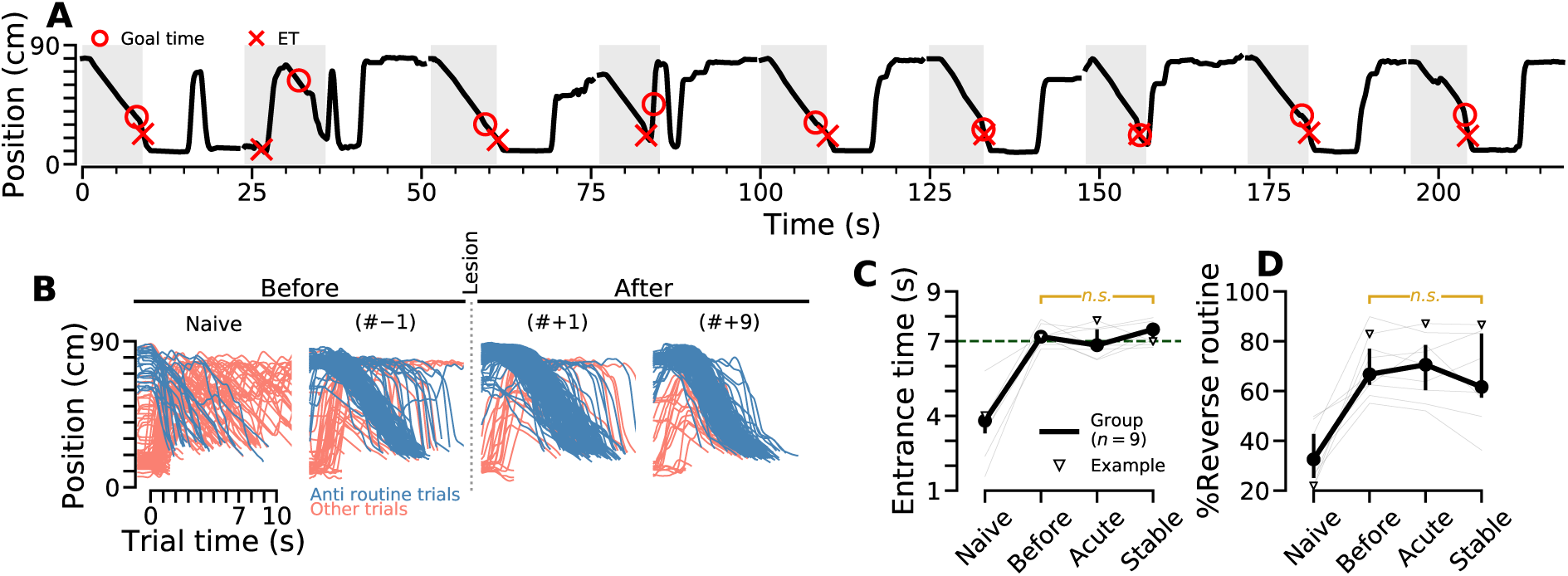
Preserved performance of a run-and-wait routine following striatal lesion. **(A)** Trajectory of a proficient animal trained in a version of the timing task in which the belt moved toward the reward area (rather than away from it). 9 consecutive trials (shaded areas) and intertrials (white areas) are shown. **(B)** Trajectories from a single representative animal in two sessions before and two sessions after lesion. **(C-D)** Comparison of *ET* (C) and percentage of run-and-wait (reverse) routine usage (D), before and after striatal lesion.

Altogether, these results indicate that the dS is not required to initiate or execute the sequential steps of a learned motor routine but is critical to invigorate its reward-oriented running phase. To better understand the origin of this vigor deficit, we examined whether the dS lesions affected the animals’ elementary ability to move. First, we compared basic locomotor activity between non-lesioned (control) and lesioned rats, in a paradigm that did not include reward-oriented runs, using a different treadmill. The locomotion test consisted of several trials (30 s long) at fixed speeds (0 to 40 cm/s), interleaved with 30 s long intertrial pauses. We found that both control and lesioned rats displayed similar exploratory locomotor activity when the treadmill remained immobile (Fig. 3A). In addition, in trials in which the treadmill was turned on, both groups were similarly able to follow a reasonable range of imposed speeds even though, as the speed increased, animals with a dS lesion ran with slightly slower speeds than control animals (Fig. 3B). It has been previously shown that the speed of reward-oriented movements increases with movement distance, to minimize temporal discounting of rewards [18, 19]. We compared running speed in trials during which running epochs were initiated from the back versus the middle of the treadmill (Fig. 3C, i.e., long vs short run distance). As predicted, animals ran faster when they initiated their runs from the back of the treadmill than from its middle (Fig. 3D). This modulation was maintained after dS lesion (Fig. 3D), although running speeds were generally slower following dS lesion. Altogether, this set of experiments indicated that the dS lesion spared basic motor abilities.

**Figure 3:**
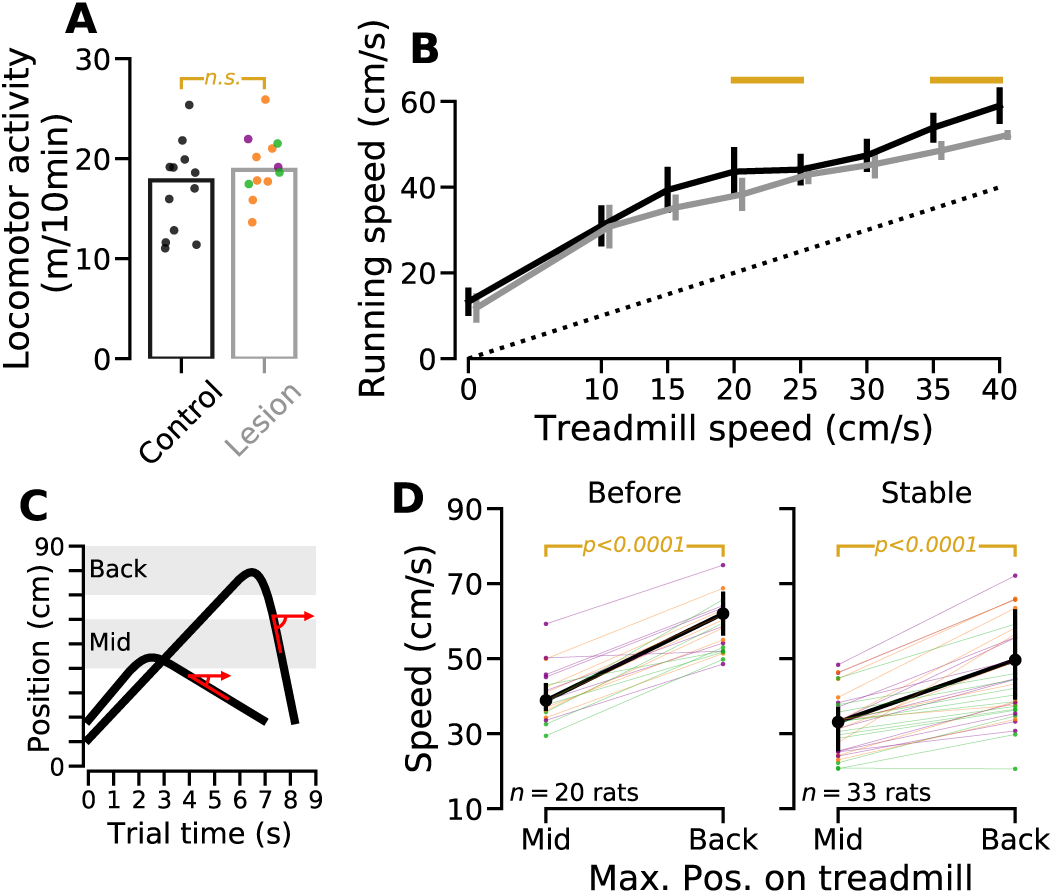
Preserved spontaneous locomotor activity and modulation of running speed following striatal lesion. **(A)** Distance ran while exploring a new (and immobile) treadmill for non-lesioned (control) and lesioned rats (*n* = 12, same color code for individual lesion type as in Fig. 1). **(B)** Average running speed in a free running task (no reward) in which control and lesioned rats were submitted to trials with incremental treadmill speed (same color code as in A). Golden lines indicated significant differences between groups (corrected for multiple comparisons). **(C)** Trials were split into 2 categories depending on whether rats initiated their run from the middle or back portion of the treadmill. Speed was computed and averaged across trial type. **(D)** Speed of the runs initiated from either the middle or back portion of the treadmill, and calculated for each animal over the last 5 sessions before lesion (left) and the last 5 stable sessions after lesion (right).

Given that the dS lesion spared the rats’ ability to modulate their running speed, the most parsimonious explanation of our results is that lesioned animals implicitly “preferred” slower speeds. A similar conclusion has been previously reached when studying the origin of bradykinesia in Parkinson’s disease (PD), which led to the proposal that such reluctance to perform fast movements was caused by an increased sensitivity to movement cost (or effort) [20]. Could a similar interpretation apply to our results? To address this question, we took advantage of the optimal control framework that relies on the assumption that animal behavior is optimal with respect to a cost function [21]. We modeled the optimal trajectory of a rat, taking into account costs related to locomotion control (effort) and those imposed by the task rules (running in the front is costly as this leads to premature *ET*, which is penalized). The effort-related term *e*(*t*) had a quadratic dependence either on the instantaneous speed or on the instantaneous muscular force produced by the animal at each time *t* during a trial. The “spatial” cost dictated by the task rules *p*(*x*) penalized positions *x* close to the reward area and was either localized or diffuse (to take into account that rats may or may not be aware of the exact location of the reward area). The aforementioned approximations of effort-related and spatial costs were combined to create for 4 versions of the model. Importantly, we computed the optimal trajectory in fixed time *T* (= 7 s) with known initial/final positions. This was done because, after lesion, rats kept initiating trials in the reward area and approached it close to the GT. Thus, in a trial of duration *T*, we assumed that the rats minimized the total cost *C*, which was a linear combination of the effort and spatial penalty terms:

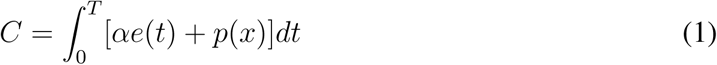

The parameter *α* governed the effort sensitivity. Fig. 4A shows how the optimal trajectories were affected by the effort sensitivity in the 4 versions of the model. In all the versions, increasing effort sensitivity (i.e., *α*) consistently resulted in optimal trajectories that laid closer and closer to the reward area, hence slow speeds were favored (Fig. 4A). We noted that this effect was more pronounced when the spatial cost was diffuse (Fig. 4A, second and fourth panels). This result is not only in agreement with the reduced running speed observed following lesion but is also reminiscent of the behavior observed during the first post-lesion session when animals with a large dS lesion ran mainly close to the reward area (Fig 1, F and G, Fig. S7). We thus reanalyzed the effect of dS lesion, focusing on animals for which the lesion caused a significant reduction in running speed (Fig. 4B, i.e., animals with larger lesions) and on trials during which animals perfectly executed the wait-and-run routine. Following dS lesion, rats started running forward earlier relative to the length of the treadmill (Fig. 4C). This effect, while being maximal on the first post-lesion sessions, remained significant after three weeks of daily testing (Fig. 4D) and was correlated with lesion size (Fig. 4E).

**Figure 4:**
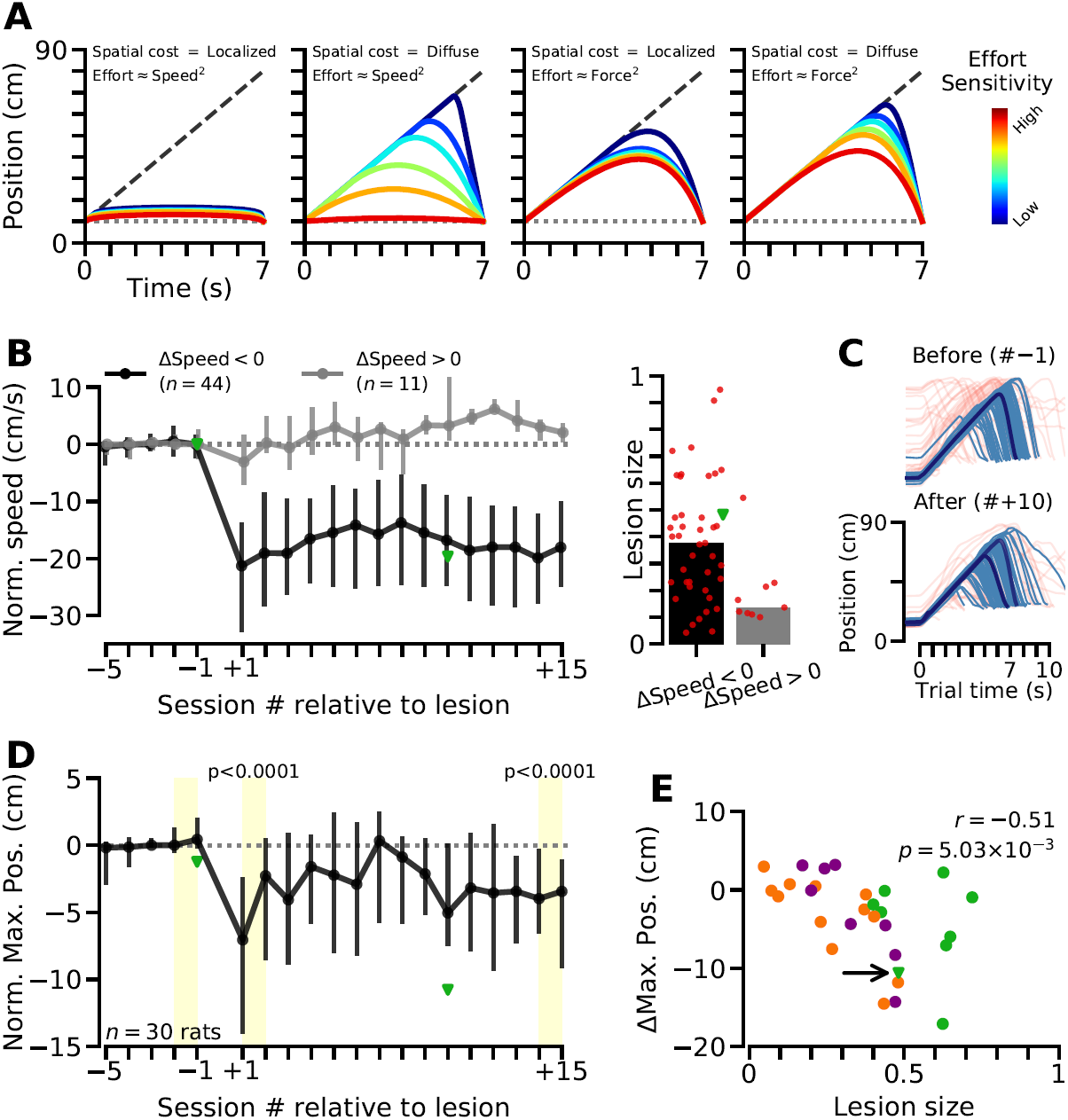
Optimal trajectory vs effort sensibility and experimental validation. **(A)** Optimal trajectories predicted by models with different effort and spatial costs approximations. The cost of premature entrance in the reward area (spatial cost) was simulated using a Heaviside function that was either localized (∼ step function with non-zero value within the reward area) or diffuse (∼ a sigmoid function whose value gradually decreases toward zero away from the reward area). Effort was approximated as the square value of either the muscular force produced by the animals or of its speed. **(B)** Left, animals were divided into two groups based on the significance of the dS lesion effect on running speed. Right, lesion size for animals in those two groups. **(C)** Effect of dS lesion on the trajectories of a single animal. Only trials in which the routine was executed (thin blue lines) were taken into account to find the trajectory of the trial with the median maximum position (bold blue line). **(D)** Effect of dS lesion on the maximum position of routine trials. **(E)** Effect of dS lesion on the maximum position versus lesion size. Same color code for individual lesion type as in Fig 1. Green triangles in B, D and E are data points from the example animal whose trajectories, before and after lesion, are shown in C.

Our results support the view that the dS lesion increased the animals’ sensitivity to effort which led them to modify how they performed the wait-and-run routine (kinematics changes). Remaining very close to the reward area minimizes energy expenditure (effort) by limiting the usage of fast speeds (Fig. 4A). This strategy was observed particularly in animals with a larger lesion (DS group in Fig. 1H and Fig. S7) but led to premature entrances in the reward area and an abrupt drop in correct trials. Noticeably, most rats with a DS lesion (14 out of 16) progressively waited longer in the following post-lesion sessions and recovered task proficiency after a few sessions (Fig. 1H). Thus, the dS may not be critical to learn the wait-and-run routine. Accordingly, we found that all the rats with DLS or DMS lesions performed before training, and the majority of the animals with DS lesions, learned our waiting task similarly than non-lesioned rats (Fig. S8A). Still, a few rats with the largest lesions (DS lesion before training, 6 animals marked in Fig. S8 and with mean lesion size of 75%; DS lesion after training, 2 rats with lesion size>80%, Fig. 1H, right, two lines crossing x-axis) ran mostly in the reward area as long as we trained them after the lesion, a strategy that may be considered as an attempt to limit energy expenditure due to high sensitivity to effort (see Fig. 4A). Thus, very large dS lesions performed before and after training, by increasing massively effort sensitivity, may irreversibly interfere with learning and performing the wait-and-run routine, respectively. Still, a robust negative correlation between running speed and lesion size was observed when we pooled data from animals with dS lesions performed before and after training (*n* = 116 animals, Fig. S4B). In addition, comparing the average maximal position of the trajectories of all control and lesioned animals showed that those with dS lesions initiated earlier (in space and time) their runs toward the reward area (Fig. S9).

Overall, our results indicate that lesioning the dorsal striatum of rats affected their performance of a motor routine in a way that is most parsimoniously explained by an increased sensitivity to effort (movement energy cost). After dS lesion, animals kept arriving on time in the reward area, but they started to run earlier (i.e., on a more intermediate portion of the treadmill) and used slower speeds to cross the treadmill. The same effect would have been expected had we forced non-lesioned rats to perform the task with extra weight on their back [19]. A similar increase in sensitivity to movement energy cost has been proposed to explain bradykinesia in Parkinson’s disease (PD) patients [20], which has led to the suggestion that the basal ganglia estimate the “cost-to-go” during execution of a motor task [22, 23]. On the one hand, there is evidence from the MitoPark mice model of PD that dopamine in the dS is necessary for the invigoration of voluntary actions [24]. On the other hand, whether the dorsal basal ganglia circuit is critical for procedural memory retrieval, and more generally for action selection, has been an important topic of debate [25, 26]. In this context, our study provides compelling evidence for a specific role of the dS in setting the sensitivity to effort, rather than in the storage or retrieval of procedural memory. Our results indicate that dS lesions changed the kinematics of a well-learned motor routine as a result of increased sensitivity to effort, without altering the animals’ capacity to run at different speeds. The selective perturbation of the activity of dS projection neurons bidirectionally modulates the speed of execution of purposive movements [15]. Our results complement this study by proposing that dS projection neurons bidirectionally modulate the perception of the effort necessary to execute a movement at a given speed, which in turn influences whether this speed will be selected next time this movement will be performed. Importantly, a contribution of the dS in the cost/benefit analysis of actions [27] has repercussions beyond the control of their execution speed and can also explain why manipulations of striatal activity change the preference for certain actions or behavioral states [2, 28, 12, 29], bias decision making [30] or alter procedural skills [31, 9, 10].

Expending effort to produce faster movements allows limiting the temporal discounting of reward (i.e., cost of time, [19]). In sensory guided decision-making tasks, the cost of time can also be reduced by limiting the duration of deliberation [32]. Interestingly, recent evidence supports a role of the basal ganglia in signaling the urgency to commit to a choice [33]. Future studies should investigate whether signaling effort and urgency are the two sides of a unique function implemented in the basal ganglia to maximize the reward rate while minimizing costs.

## Acknowledgements

We thank Drs. I. Bureau, R. Cossart, J. Epsztein, E. Fino, J. de la Rocha and T. Verstynen for critical reading of the manuscript and Dr A. de Chevigny for advice with histological verification of the lesions.

## Author contributions

Maria-Teresa Jurado-Parras: Conceptualization, Investigation, Writing–review & editing; Mostafa Safaie: Conceptualization, Software, Visualization, Investigation, Formal analysis, Writing–original draft; Stefania Sarno: Software, Methodology, Formal analysis, Conceptualization, Visualization; Jordane Louis: Investigation; Corane Karoutchi: Investigation; Bastien Berret: Conceptualization, Methodology, Formal analysis, Writing–review & editing; David Robbe: Funding acquisition, Project administration, Supervision, Visualization, Writing–original draft, Conceptualization, Software.

## Supplementary materials

### Supplementary Figures

**Supplementary Figure S1:**
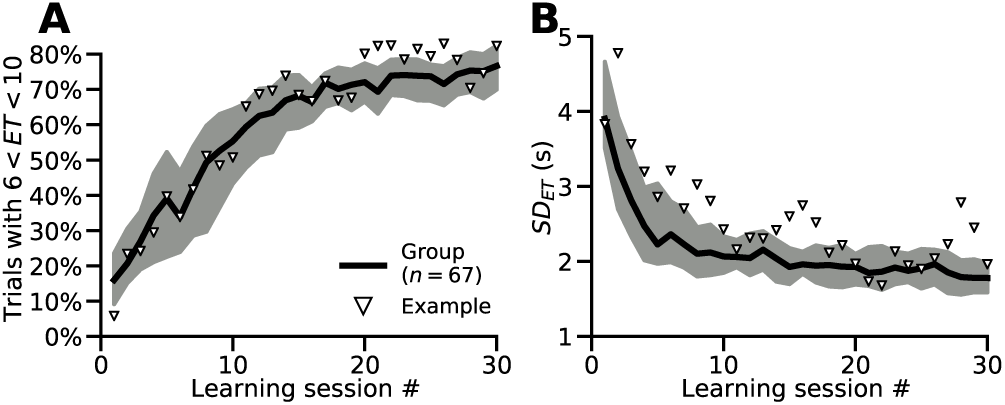
Task performance improvement across sessions. **(A)** Percentage of trials in which animals entered the reward area close to *GT* (6 *s* < *ET* < 10 *s*), session-by-session. **(B)** Session-by-session standard deviation of *ET*. Triangles show performance improvement for an example animal (same animal as in Fig. 1).

**Supplementary Figure S2:**
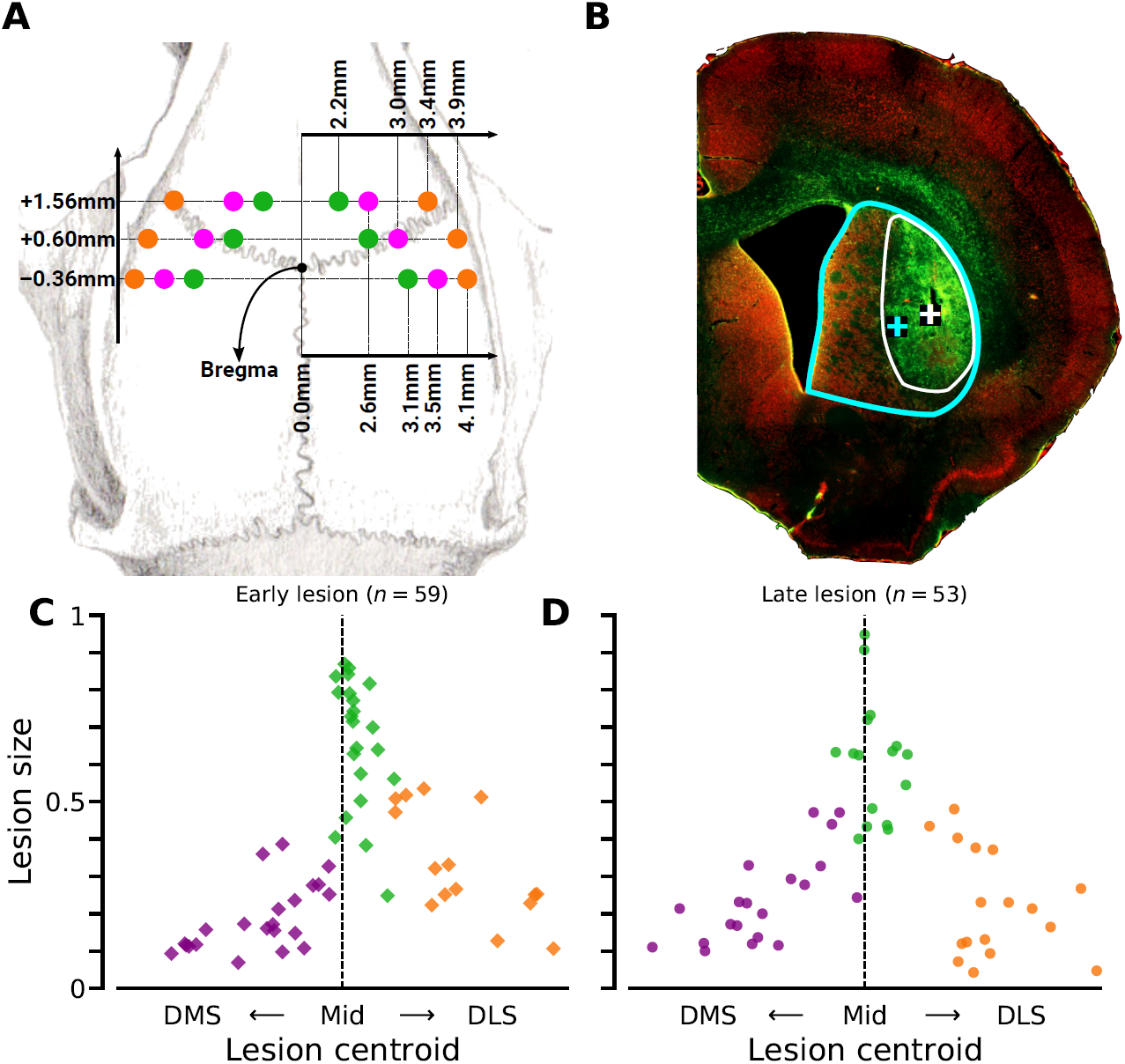
Dorsal striatum lesion quantification. **(A)** Schematic of the lesion sites. **(B)** Illustration of the quantification of the lesion size. For each coronal slide and hemi-striatum, the contour of the lesion was manually outlined using GFAP staining. The relative size of the lesion (compared to the full dS, manually outlined on the NeuN staining) and the coordinates of the lesion/striatum centroid was calculated. For each animal, the size and laterality were obtained by averaging data along the anteroposterior axis, for both left and right hemispheres. **(C, D)** Lesion size versus laterality for animals that underwent lesion before (Early, C) and after (Late, D) extensive practice. Lesion quantification was performed blindly relative to behavioral analysis. In four animals with a dS lesion performed after learning the task (late lesion), the lesion size quantification could not be properly performed. These animals were classified according to their injection coordinates in the surgery (3 DLS and 1 DMS), however they were excluded from any analyses that required the lesion size (hence the difference between the number of “late lesion” animals in this figure, *n* = 53, and the total number of animals in Fig. 1, *n* = 57).

**Supplementary Figure S3:**
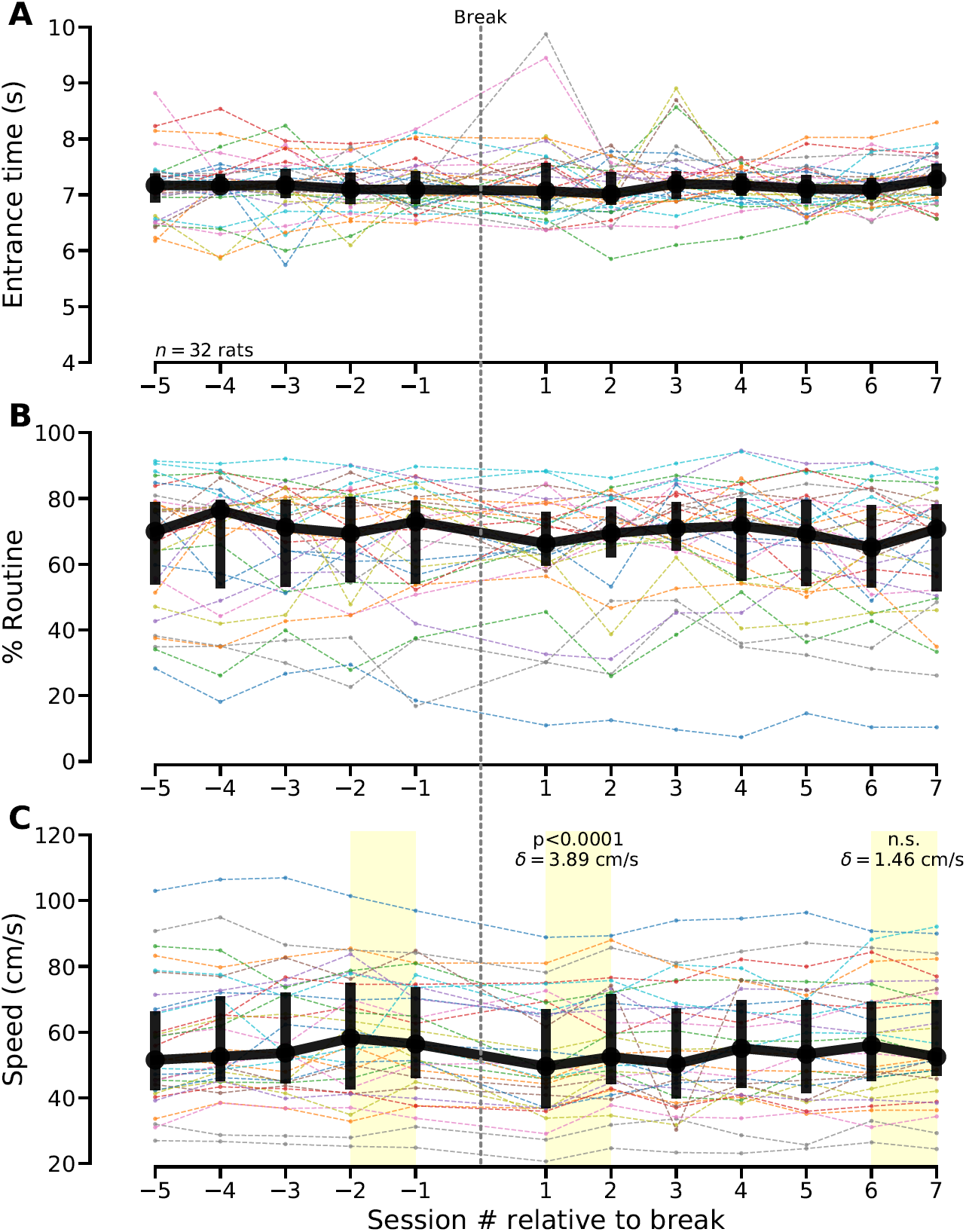
Impact of a two weeks break in practice on performance. **(A-C)** Task performance before and after a two weeks-long break in practice. Non-lesioned animals had stable performance before the two weeks-long break (same duration than lesion recovery period) in practice. (A), Entrance time. (B) Percentage of trials during which animals used the wait-and-run routine. (C) Speed of the animals when they ran toward the reward area. A small but significant reduction in running speed was observed just after the break (*δ* denotes the effect size).

**Supplementary Figure S4:**
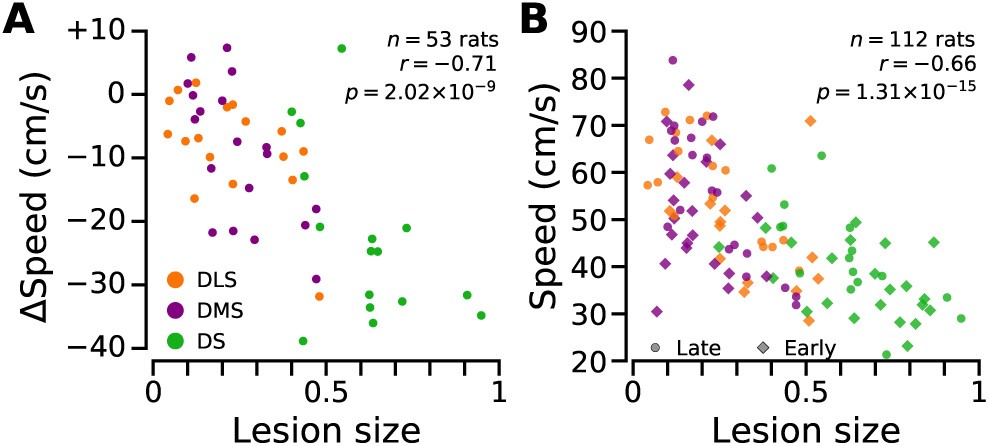
The impact of the dS lesion on running speed correlates with lesion size. **(A)** Average change in running speed (speed after lesion − speed before lesion) versus lesion size, for all the rats that received a striatal lesion after training (late lesion). Running speed was calculated when rats crossed the treadmill from its back region to the reward area. All the running speed values obtained across 5 consecutive sessions were averaged to obtain the average running speed before (last 5 sessions before lesion break) and after (sessions #4 to #9, relative to lesion break) lesion. **(B)** Average running speed versus lesion size for all the animals that underwent surgical lesion of the dS. This dataset (*n* = 112 animals) includes all the animals that underwent striatal lesion (DLS, DMS, DS) after extensive training (Late group, *n* = 53, same animals as in panel A), and animals that underwent lesion before training (Early group, *n* = 59 animals). Speed was computed as in A, except that average was done over the last 5 sessions (for both Early and Late groups). In four animals with a dS lesion performed after learning the task (late lesion), the lesion size quantification could not be properly performed. These animals were classified according to their injection coordinates in the surgery (3 DLS and 1 DMS), however they were excluded from any analyses that required the lesion size (hence the difference between the number of “late lesion” animals in this figure, *n* = 53, and the total number of animals in Fig. 1, *n* = 57).

**Supplementary Figure S5:**
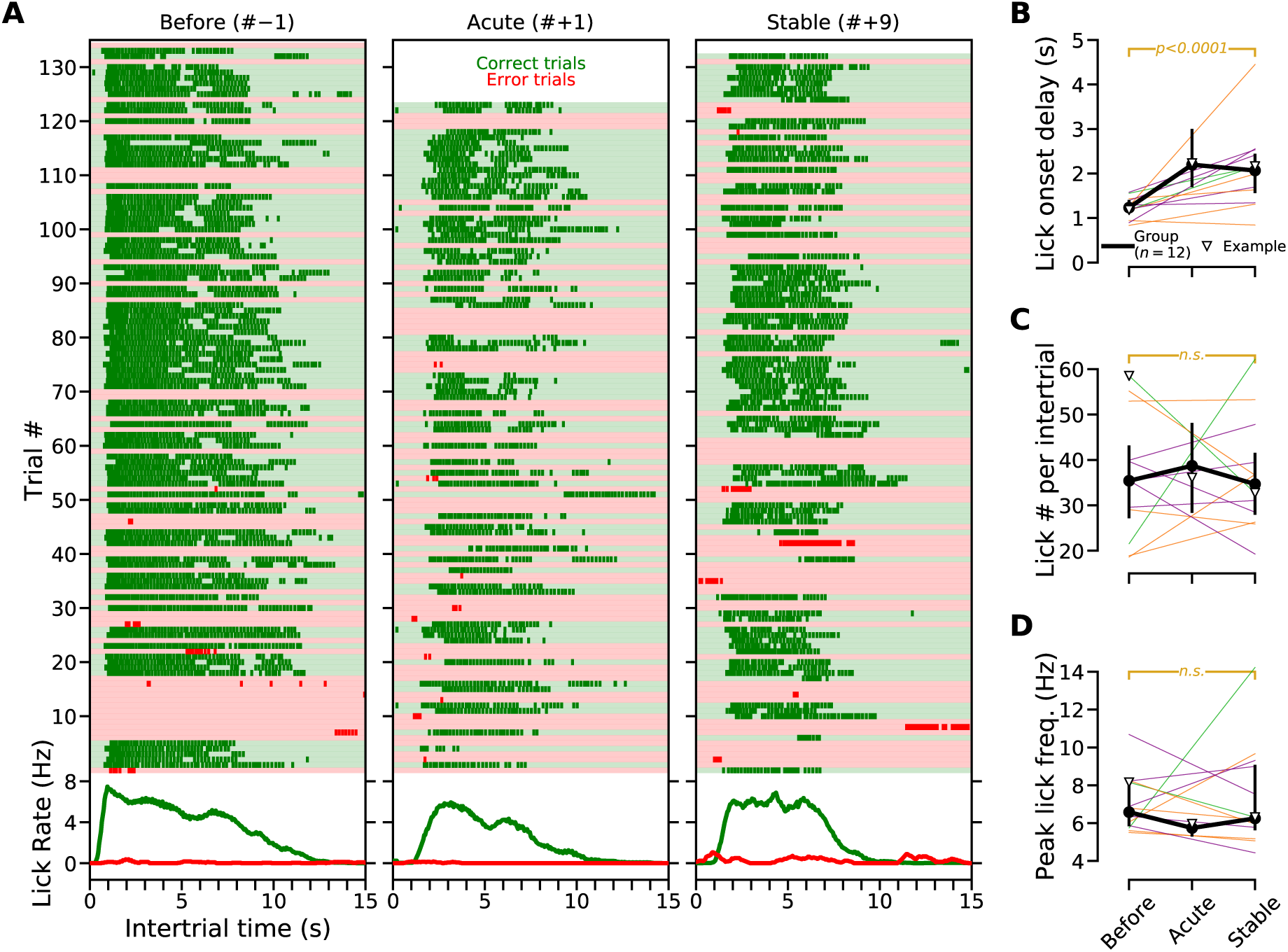
Licking behavior after dS lesion. **(A)** Trial-by-trial licking pattern (top) and averaged lick rate aligned to intertrial onset for a single animal in 3 sessions (1 just before and 2 after lesion). **(B-D)** Effect of dS lesion on lick onset delay (B), number of licks per intertrial (C) and peak lick frequency (D) Same color code for individual lesion type as in Fig. 1 or Fig. S4.

**Supplementary Figure S6:**
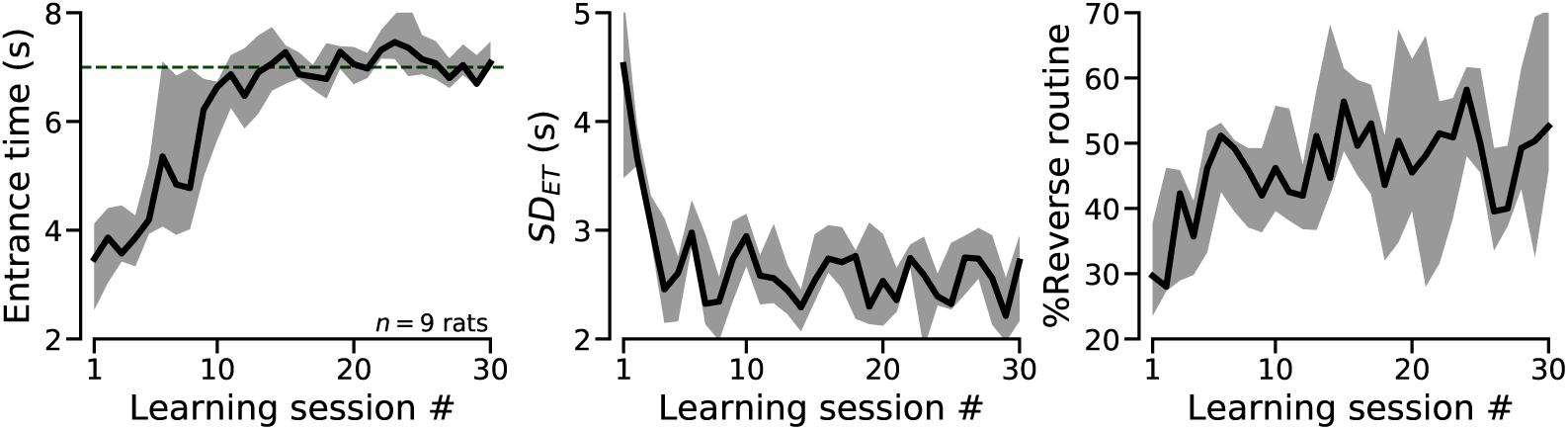
Performance improvement in the reverse treadmill task. Left: Entrance time across learning sessions. Middle: Session-by-session standard deviation of *ET*. Right: Percentage of trials during which animals used the run-and-wait (reverse) routine.

**Supplementary Figure S7:**
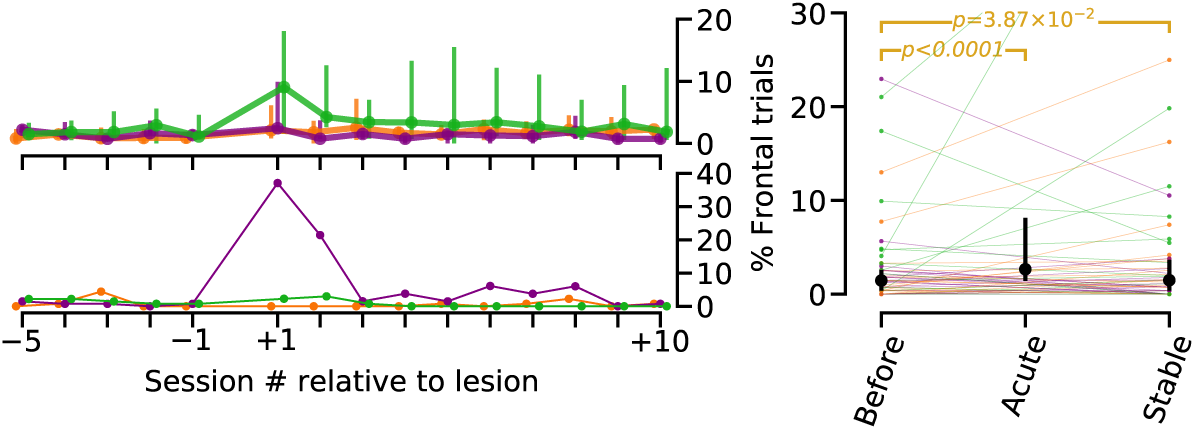
Dorsal striatum lesion induced a transient increase in the percentage of trials during which animals remained close to the reward area. Upper left: Session-by-session percentage of trials in which animals remained close to the reward area (Frontal trials). Group data for animals with DLS, DMS and DS lesion. Same color code as in Fig. 1. Lower left: Same as above, but for the illustrative animals shown in Fig. 1, E to F. Right: Statistical comparisons before and after dS lesion. Same color code for individual lesion type as in Fig. 1, Fig. S4.

**Supplementary Figure S8:**
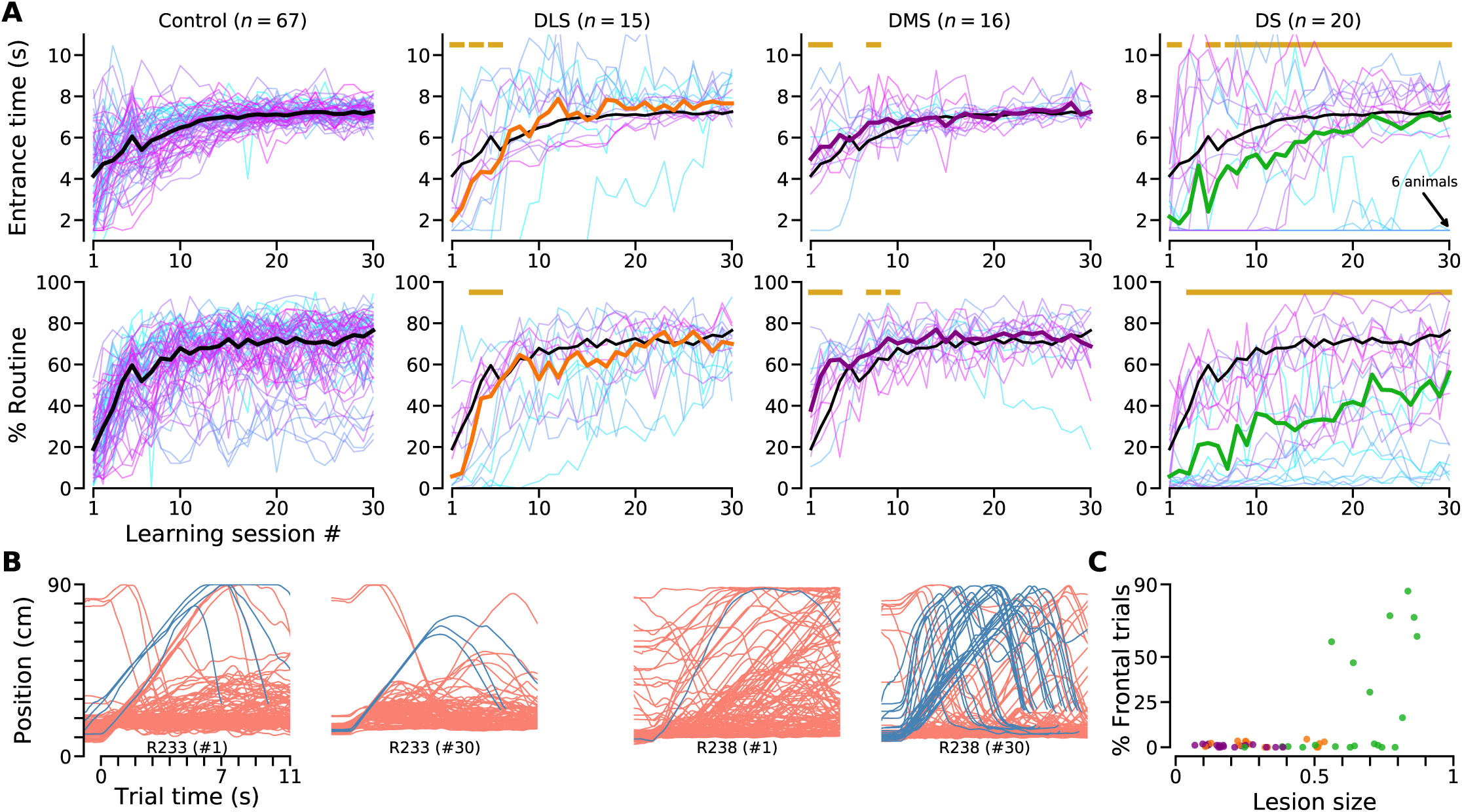
Effect of DLS, DMS and DS lesions performed before training on task learning. **(A)** Session-by-session change in performance (*ET*, upper panels; Percentage of trials in which the routine was used, lower panels) for animals without lesion (Control, left) and for animals that received a lesion before training (DLS, DMS, DS from left to right). Black lines indicate Control group median. Thin colored lines indicate single animals. Thick colored lines (same color code as in Fig. 1) in 3 rightmost columns indicate group performance for comparison (8 lesion animals with fewer than 30 training sessions are not shown, which explains the difference in the number of animals in this figure and fig. S2C). Horizontal golden lines indicate significant differences between control and lesion groups (corrected for multiple comparisons). **(B)** Trajectories before and after extensive training (sessions #1 and #30) for two animals with large DS lesions. Note that, after extensive practice, R238 was capable of performing the wait-and-run routine. **(C)** Percentage of trials in which animals remained in the front region of the treadmill (computed for sessions #25 to #30) versus lesion size.

**Supplementary Figure S9:**
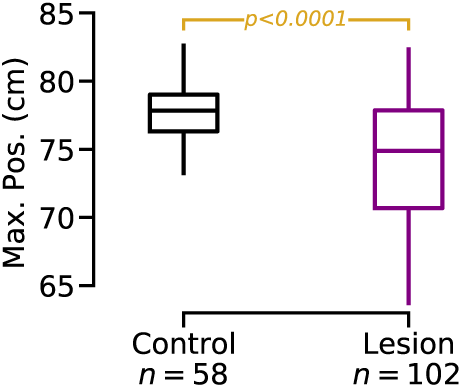
Maximum position of the average trajectory of control and lesioned rats. Each boxplot represents the range of the Max. Pos. (center line, median; box, 25th and 75th percentiles; whiskers, 5th and 95th percentiles) for control and lesioned (before and after training) animals. For each animal, the median value of their average trajectory was computed over the last 5 sessions performed without lesion (Control group, at least 30 training sessions), and/or the last 5 sessions performed after dS lesion (Lesion group). The difference in number of control animals between this figure (*n* = 58) and Fig. S8 (*n* = 67) is due to 9 animals that did not perform the wait-and-run routine reliably, precluding the computation of the Max. Pos. value. Comparison using the permutation test (10000 permutations).

## Materials and Methods

### Subjects

A total of 166 male Long-Evans rats were used in this study (number for each experimental condition is directly reported in the figures). They were 12 weeks old at the beginning of the experiments, housed in groups of 4 rats in temperature-controlled ventilated racks and kept under 12 h–12 h light/dark cycle. All the experiments were performed during the light cycle. Food was available *ad libitum* in their homecage. Rats had restricted access to water while their body weights were regularly measured. No animal was manually excluded from the analysis. All experimental procedures were conducted in accordance with standard ethical guidelines (European Communities Directive 86/60 - EEC) and were approved by the relevant national ethics committee (Ministère de l’enseignement supérieur et de la recherche, France, Authorizations #00172.01 and #16195).

### Apparatus

Four identical treadmills were used for the experiments. Each treadmill was placed inside a sound-attenuating box. Treadmills were 90 cm long and 14 cm wide, surrounded by plexiglass walls such that the animals were completely confined on top of the treadmill conveyor belt. The treadmill belt covered the entire floor surface and was driven by a brushless digital motor (BGB 44 SI, Dunkermotoren). The front wall (relative to the turning direction of the belt) was equipped with a device delivering drops of 10% sucrose water solution (maximal drop size ∼ 80 *µ*L). An infrared beam, located at 10 cm of this device, defined the limit of the reward area. A loudspeaker placed outside the treadmill was used to play an auditory noise (1.5 kHz, 65 db) to signal incorrect behavior (see below). Two strips of LED lights were installed on the ceiling along the treadmill to provide visible and infrared lighting during trials and intertrials, respectively (see below). The animals’ position was tracked via a ceiling-mounted camera (Basler scout, 25 fps). A custom-made algorithm detected the animal’s body and recorded its centroid to approximate its position on the treadmill. After trial onset, the first interruption of the beam was registered as entrance time in the reward area (*ET*). The entire setup was fully automated by a custom-made program (LabVIEW, National Instruments). The experimenter was never present in the behavioral laboratory during the experiments.

### Habituation

Animals were handled 30 m per day for 3 days, then habituated to the treadmill for 3 to 5 daily sessions of 30 min, while the treadmill’s motor remained turned off and a drop of sucrose solution was delivered every minute. Habituation sessions resulted in systematic consumption of the reward upon delivery and a tendency of the animals to spend more time in the reward area.

### Main Behavioral Task

Training started after handling and habituation. Each animal was trained once a day, 5 times a week (no training on weekends). Each of the daily sessions lasted for 55 min and contained ∼ 130 trials. Trials were separated by intertrial periods of 15 s. During intertrials, the treadmill remained dark and infrared ceiling-mounted LEDs were turned on to enable video tracking of the animals. Position was not recorded during the last second of the intertrials to avoid buffer overflow of our tracking routine and allow for writing to the disk (see the gaps in the position trace in Fig. 2A). The beginning of each trial was cued by turning on the ambient light, 1 s before motor onset. Since animals developed a preference for the reward area during habituation, the infrared beam was turned on 1.5 s after trial start. This delay was sufficient to let the animals be carried out of the reward area by the treadmill, provided they did not move forward. After the first 1.5 s, the first interruption of the beam was considered as *ET*. The outcome of the trial depended solely on the value of the *ET*, compared to the goal time (GT= 7 s). In a correct trial (*ET* ≥ *GT*), an infrared beam crossing stopped the motor, turned off the ambient light, and triggered the delivery of reward. In an error trial (*ET* < *GT*), there was an extended running penalty. During the penalty, the motor kept running, the ambient light stayed on and an audio noise indicated an error trial. The duration of the penalty period was anticorrelated with the magnitude of error, between 1 s and 10 s (see [1] for more details). In trials wherein animals didn’t cross the beam in 15 s since trial onset, trial stopped and reward was not delivered.

The magnitude of the reward was a function of the *ET* and animal’s performance in previous sessions (only in early training). Reward was maximal at *ET* = *GT* and dropped linearly to a minimum (= 38% of the maximum) for *ET* s approaching 15 s (i.e., the maximum trial duration). Moreover, in the beginning of the training, partial reward was also delivered for error trials with *ET* > *ET*_0_, where *ET*_0_ denotes the minimum threshold for getting a reward. The magnitude of this additional reward increased linearly from zero for *ET* = *ET*_0_, to its maximum volume for *ET* = *GT*. In the first session of training, *ET*_0_ = 1.5 s and for every following session, it was updated to the maximum value of median *ET* s of the past sessions. Once *ET*_0_ reached the *GT*, it was not updated anymore.

### Reverse Treadmill Task

This task differed from the normal task in three critical properties: 1) the treadmill moving direction was reversed, i.e., the conveyor belt moved toward the reward port; 2) the treadmill speed was set at 8 cm/s (instead of 10 cm/s) to ensure that starting the trial in the back of the treadmill and remaining still after trial onset would be rewarded, i.e., *ET* ≥ 7. 3) the intertrial duration was 20 s, instead of 15 s, to allow sufficient time for the animals to move to the back of the treadmill while the motor was still off.

### Locomotor Activity Test

A group of animals with a striatal lesion (7 DLS, 2 DMS, and 3 DS), and another group of non-lesioned animals (*n* = 12) were used in this test to assess their general locomotor activity. Prior to this task, animals had full access to water and food for at least 3 days. Then, they were placed on an unfamiliar treadmill, with a different structure (slanted walls and covered reward port) compared to the treadmill in which they were trained, while their position was being recorded using a side-mounted high-speed camera (200 fps). During the first 10 min, the ambient light was turned off and the treadmill remained immobile. Their exploratory locomotor activity, i.e., how much they moved along the treadmill, during this period is presented in Fig. 3C. Then, in a free-running task, they ran in trials of 30 s while the treadmill speed progressively increased across trials (5 trials at 0 cm/s, 2 trials at 10 cm/s, 3 trials at 15 cm/s and 5 trials at 20, 25, 30, 35, and 40 cm/s, data shown in Fig. 3D). Each trial was followed by an intertrial (30 s-long), with the ambient light and the treadmill motor were turned off. The running speed reported in Fig. 3D is the average running speed of animals during the trials of any given treadmill speed.

### Lesion Surgery

Anesthesia was induced with an intraperitoneal (IP) injection of a mixture of 100 mg/kg ketamine and 10 mg/kg xylazine and was maintained during the surgery with inhalant isoflurane gas (less than 3%). After shaving and cleaning the scalp, the animal was placed in the stereotaxic frame (Kopf Instruments) and a local anesthetic (lidocaine) was injected under the scalp. Then, an incision along the midline of the skull was made, allowing for cleaning the exposed skull and drilling the craniotomies above the targeted areas. To perform fiber-sparing lesion of the striatum, ibotenic acid (1% in 0.1M NaOH, Fisher Scientific) was infused (Pump 11 Elite Nanomite, Harvard Apparatus, using a 10 uL WPI Nanofil syringe) in 6 specular sites bilaterally, at a rate of 90 nl/min. Injection coordinates (in mm, with reference to Bregma, according to Paxinos) are shown in Fig. S2A (each injection at −5.6 dorsoventral). The infused volume in each site was 200 nL for DLS and DMS lesions, and 400 nL for DS lesions. The needle remained in place for 10 min following the injection to allow for the diffusion of the excitotoxin. Then the needle was retracted slowly to avoid backflow of the drug. Once all the injections were performed, craniotomies were filled with bone wax, the skull was disinfected and the skin was sutured. Animals were allowed to recover for two weeks before resuming behavioral training. After surgery, animals were housed alone for 3 days in a warmed cage, to avoid getting hurt by their cagemates, and were force-fed if needed. Post-surgery pain was reduced as much as possible using an opiate painkiller (Buprenorphine) and if necessary a non-steroidal anti-inflammatory drug (Carprofen).

### Histology

Animals were euthanized with an overdose IP injection of 2 mL pentobarbital or with an injection of Zoletil (40mg/kg) and Domitor (0.6 mg/kg). Then, they were perfused with 4% paraformaldehyde and their brains were harvested for histological analysis of the lesion size and location. Brains were coronally sliced on a vibratome at 60 um thickness. For each animal, six sections spanning the dorsal striatum along the rostrocaudal axis were selected (usually the following slice numbers: 5, 15, 25, 35, 45, and 55 for consistency) and submerged in 0.1 M PBS. Then, PBS was replaced with citrate buffer (10 mM) for 10 min at room temperature. Next, slices were submerged with a blocking solution, consisting of PBS with 0.3% Triton and 15% normal goat serum (NGS) for 2 hrs at room temperature. Then the solution was replaced with a solution consisting of 2 uL anti-NeuN antibody (Merks Millipore, MAB377) and 0.5 uL of anti-GFAP antibody (Agilent, Z033429-2) diluted in 200 uL of the blocking solution, kept overnight at 4° C. Sections were then rinsed twice in PBS for 10 min at room temperature, before being resubmerged in 1 uL of donkey-anti-mouse antibody (Al555, red), 1 uL of donkey-anti-rabbit antibody (AL488, green) diluted in 400 uL of PBS for 2 hr at room temperature. Finally, they were washed twice in PBS for 10 min and mounted for microscopy.

### Data Analysis

Data from each behavioral session was stored in separate text files, containing position information, entrance times, treadmill speeds, and all the task parameters. Position information was then smoothed (Gaussian kernel, *σ* = 0.3 s). The entire data processing pipeline was implemented in python, using open-source libraries and custom-made scripts. We used a series of Jupyter Notebooks to process, quantify, and visualize every aspect of behavior, to develop and run the optimal control simulations, and to generate all the figures in this manuscript. The Jupyter Notebooks and the raw data necessary for the full replication of the figures will be publicly available via the Open Science Foundation, as done in our two last manuscripts [2, 1].

#### Motor Routine Definition

We quantified the percentage of trials in which animals performed the wait-and-run motor routine in each session (Fig. 1 I). A trial was considered *routine* if all the following three conditions were met: 1) the animal started the trial in the front (initial position < 30*cm*); 2) the animal reached the rear portion of the treadmill during the trial (maximum trial position > 50*cm*); 3) the animal completed the trial (i.e., it crossed the infrared beam).

#### Speed Calculation

Unless otherwise stated, speed in this manuscript refers to the velocity with which animals crossed the treadmill toward the reward port. For every trial, it is calculated based on the time the animal takes to run from 60 cm to 40 cm along the treadmill. Speed for each training session is the average speed across its trials (Fig. 1 J). Furthermore, in Fig. 4B, we categorized the animals based on whether they had an effect on their running speed after dS lesion (black), or not (gray). Animals were assigned to the black group (ΔSpeed< 0) if the average speed of 5 consecutive stable sessions after the lesion (i.e., session +8 to +13) were lower than that of 5 consecutive sessions before the surgery (i.e., sessions −5 to −1).

#### Reverse Routine Definition

Percentage of reverse (run-and-wait instead of wait-and-run) routine trials is analogous to the percentage of routine trials, except that it is performed in the version of the task in which the treadmill belt is moving toward the reward. A trial was considered *reverse routine* if the following conditions were met: 1) the animal started the trial in the back region of the treadmill(initial position > 60*cm*); 2) the animal completed the trial (i.e., it crossed the infrared beam).

#### Definition of Frontal Trials

Frontal trials are defined as trials in which the animal remained in the frontal portion of the treadmill (i.e., position < 30 cm) for the first 5 s after trial onset.

#### Speed Modulation Analysis

In Fig. 3D, we split the trajectories that strictly followed the wait-and-run routine (see the definition of the Max. Pos.) into trials with the maximum position between 40 and 60 cm (Mid) and those between 70 and 90 cm (Back). The data were pooled from the last 5 sessions before (Fig. 3D, left) and after (Fig. 3D, right) the lesion. To improve the reliability, animals were discarded if they did not have at least 10 trials in the Mid and 10 trials in the Back condition (trials that strictly followed the wait-and-run routine, their maximum position was within the range, and for which the speed could have been defined). The fewer number of animals in the left panel was due to the fact that most animals performed the wait-and-run routine by going all the way to the rear portion of the treadmill, thus not enough Mid trials existed.

#### Definition of Max. Pos

The maximum position an animal reached along the treadmill before initiating the run epoch toward the reward in the wait-and-run routine was quantified as Max. Pos. in Fig. 4D. Therefore, Max. Pos. was only calculated for trials that strictly followed a wait-and-run routine, i.e., total immobility followed by continuous running until reaching the infrared beam. A trial was qualified if the following conditions were met: 1) the animal started the trial in the front (initial position < 30*cm*); 2) the animal moved at least 10 cm backward (maximum position ≥ 40*cm*); 3) the animal remained still while being pushed backward by the treadmill (movement shorter than 0.1 s and slower than 5 cm/s were ignored to correct for jitter in position detection); 4) the animal performed an uninterrupted running epoch (staying immobile or moving backward shorter than 0.1 s was ignored to correct for jitter in position detection); 5) the animal completed the trial (i.e., it crossed the infrared beam). Notice that compared to the definition of the routine trials, the threshold for maximum position in the second criterion is relaxed (40 cm, compared to 50 cm) to allow the detection of trials with a reduced maximum position. To increase the reliability, any session with fewer than 10 trials for which Max. Pos. could be defined was excluded from further analysis. The reported value of Max. Pos. for each session is the average value across its trials (Fig. 4D).

#### Normalizing Speed and Max. Pos

In Fig. 4B and D, to normalize each animal’s performance according to its own behavior prior to the lesion, behavioral measures (speed and Max. Pos.) of individual animals during the illustrated sessions were subtracted from the median value of the respective measure during the pre-lesion sessions. Animals were included only if the behavioral measure could be defined in at least half of the illustrated sessions. Different *n* in panel D compared to B, and B compared to the total number of animals (Fig.1H) is due to this criterion.

#### Quantification of lesion size

Whole slices were imaged using an Apotome microscope (Zeiss, 28126), and stitched together in the processing software (Zeiss Zen). Then, for each slice, ventricle, striatum, and lesioned area were manually outlined (Fig. S2B) bilaterally in the image processing software (ImageJ, Fiji). This task was performed blindly to the behavioral results. The size and the centroid coordinates were automatically computed for all of the above-mentioned areas. The anteroposterior location of each slice was also approximated according to the rat brain atlas (Paxinos). The lesion size reported in this paper is the ratio of the lesion volume over the volume of the striatum. The region of interest was approximated as a truncated cone between any two consecutive sections, and the volume was accordingly calculated. The type of lesion (DLS, DMS or DS) was also determined visually and confirmed by comparing the centroid location of the lesion to that of the entire striatum (see Fig. S2). Animals with a DLS lesion in one hemisphere and a DMS in another (*n* = 7) were excluded from this manuscript. In four animals with a dS lesion performed after learning the task, the lesion size quantification could not be properly performed. These animals were classified according to their injection coordinates in the surgery (3 DLS and 1 DMS), however, they were excluded from any analyses that required the lesion size (hence the difference between the number of animals in Fig. S4A and the total number in Fig. 1).

### Statistics

All statistical comparisons were performed using resampling methods (permutation test and bootstrapping, in every case, *n* = 10000 iterations). The permutation test used in this manuscript has already been described and implemented [1]. In brief, to compare two groups, random re-assignment of data points to surrogate groups should not generate group differences similar to that of the original groups. By repeating this process over and over and building a distribution of surrogate group differences, we estimated the likelihood of the original group difference belonging to this distribution. This test was used two compare two independent groups, controlling for multiple comparisons, in Fig. 3D, and Fig. S7. A similar permutation test was also used to compare two sets of unpaired data points, such as in Fig. 3C. For paired comparisons (in Fig. 1H-J, Fig. 2C, D, Fig. 3B, Fig. 4D, Fig. S3C and Fig. S5B-D,), we generated the bootstrap distribution of mean differences (*n* = 10000 with replacement). Significance was reported if 95% Confidence Interval (CI) of the pairwise differences differed from zero (i.e., zero was not within the CI). For example, in Fig. 1J, right, the 95% CI of pairwise differences is (−9.45, −15.25). Since this interval does not contain zero, it is reported significant, whereas in Fig. 1I, right, the CI is (−0.04, 0.03) which includes zero, and hence is reported non-significant. For significant comparisons, the p-value was then reported as the fraction of samples in the resampled distribution with their sign opposite to that of the mean. In case no samples were found, *p* < 0.0001 was reported, i.e., smaller than one chance in 10000. The relation between lesion size and magnitude of behavioral impairment was quantified using the Pearson’s correlation coefficient.

### Optimal control model description

The optimal control (OC) theory relies upon the assumption that behavior is governed by optimality principles with respect to some cost function [3]. This means that, when making a decision and generating movements, the default tendency of the brain is to maximize reward while minimizing effort. Here, we aimed at simulating the optimal trajectories of rats while they performed the wait-and-run motor sequence and when their sensitivity to effort was manipulated. Critically, because striatal lesions changed the kinematics of the motor sequence execution, we wanted to test the hypothesis that the changes in kinematics were compatible with increased sensitivity to effort. After lesion, rats kept initiating the trial in the reward area and arrived in the reward area close to the Goal Time (*GT* = 7*s*, Fig. 1). We thus computed the optimal trajectory in fixed time *T* (= 7 s) with known initial/final states, given a system dynamics and cost.

#### Equation of motion and constraints

We assumed that the speed of a rat *v* satisfies the following equation of motion[4]:

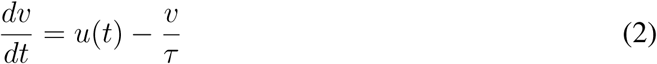

The term *v/τ* in the above equation is a resistive force per unit mass. The term *τ* is a friction coefficient (when *τ* increases resistance decreases) and *u*(*t*) is the propulsive force per unit mass. In all the simulation we set *τ* = 1.8 s. Qualitatively, the results we obtained are independent of *τ*. A key component of the behavioral task is that to obtain a reward, animals must enter the reward area after *GT*, This reward area is delimited by the front wall of the treadmill equipped with a reward port and an infrared-beam located at 10 cm of the front wall.

In agreement with the behavioral data, the initial and final positions of the animal in the model were both equal to the beam position *x*_*b*_ = 10 cm. The position of the rat is constrained by the treadmill length, which is *L*_*T*_ = 90 cm.

The treadmill pushes the animal toward the rear of the treadmill (located at position 90 cm) with a constant positive velocity of *v*_*T*_ = 10 cm/s. In the coordinate adopted in the model, running toward the reward area is, therefore, occurring with negative velocity. The velocity of the rat was also constrained to be less than 0, i.e. *v*_*max*_ = 0 cm/s (the rat cannot run backward) and its module could not be bigger than 70 cm/s, i.e. *v*_*min*_ = −70 cm/s (the rat cannot run toward the reward area at a speed faster than 70 cm/s).

We defined the components of the state vector X= [*x*^0^, *x*^1^], respectively as the position and speed of the rat in the laboratory frame of reference. Their dynamics is governed by the equations:

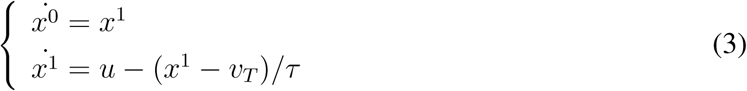

The state variables are constrained by the following initial conditions and inequalities:

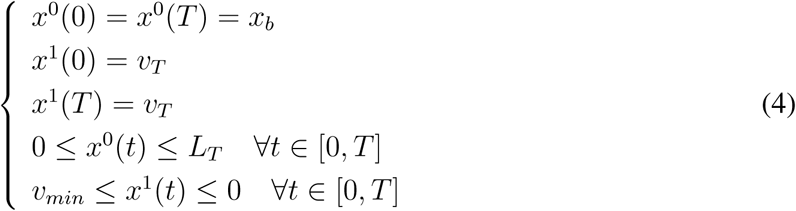

#### Control variable and cost function

In the OC framework, we assumed that a rat modulates its propulsive force on a moment-to-moment basis, so *u*(*t*) is the control. The infinitesimal energetic cost *c*(*t*) is assumed to be the linear combination of two terms. The first term is an effort-related term *e*(*t*) that has a quadratic dependence on either speed or force (i.e., on the control):

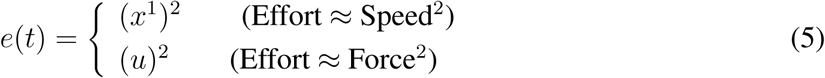

The second term is a cost related to the task rules, namely that running in the reward area before *GT* is associated with a penalty. The modeled trajectory must respect the following rule: *x*^0^(*t*) > *x*_*b*_ ∀ *t* < *T* (*x*_*b*_ being the position of the infrared beam). We modelled this “spatial” cost, using a differentiable approximation of the Heaviside function of height *A* = 10:

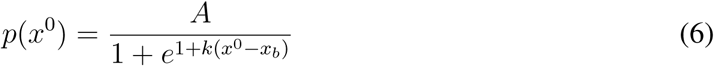

The parameter *k* governs the steepness of this spatial cost. We used *k* = 100 and *k* = 1 to model the spatial cost in a way that is either localized or diffuse, respectively (see Fig below). The rationale for doing so is that it is difficult to precisely know how the animals perceived the dangerousness of running close to the reward area.

Therefore, in a trial of duration *T*, the rat wants to minimize the total cost *C*:

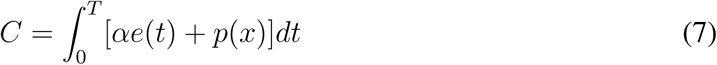

The parameter *α* governs the effort sensitivity. In the simulations, we used six values for the parameter *α*, namely *α* = {0.1, 1, 2, 5, 10, 100}. High values of *α* correspond to high effort sensitivity.

#### Numerical implementation

We used the *CasAdi* [5] software and direct collocation method to numerically find the optimal trajectories.

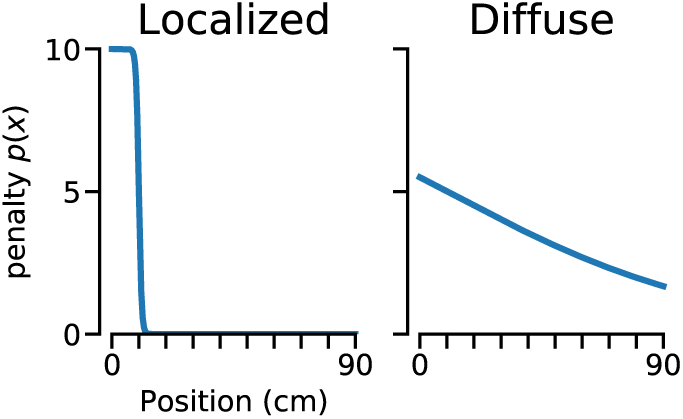

#### Penalty functions used to model the spatial cost

Localized (left) and diffuse (right) penalty functions used to model the spatial cost *p*(*x*).

